# A spectrum of clinically-identified cysteine mutations in fibulin-3 (EFEMP1) highlight its disulfide bonding complexity and potential to induce stress response activation

**DOI:** 10.1101/2024.07.22.604634

**Authors:** Gracen E. Collier, John D. Hulleman

## Abstract

Fibulin-3 (FBLN3), also known as EFEMP1, is a secreted extracellular matrix (ECM) glycoprotein that contains forty cysteine residues. These cysteines, which are distributed across one atypical and five canonical calcium-binding epidermal growth factor (EGF) domains, are important for regulating FBLN3 structure, secretion, and presumably function. As evidence of this importance, a rare homozygous p.C55R mutation in FBLN3 negates its function, alters disulfide bonding, and causes marfanoid syndrome. Additional studies suggest that heterozygous premature stop codon mutations in FBLN3 may also cause similar, albeit less severe, connective tissue disorders. Interestingly, a series of twenty-four cysteine mutations in FBLN3 have been identified in the human population and published in the Clinical Variation (ClinVar) and gnomAD databases. We tested how seven of these cysteine mutants (five loss-of-cysteine variants: C42Y, C190R, C218R, C252F, and C365S, two gain-of-cysteine variants: R358C, Y369C) and two newly developed mutations (G57C and Y397C) altered FBLN3 secretion, disulfide bonding, MMP2 zymography, and stress response activation Surprisingly, we found a wide variety of biochemical behaviors: i) loss-of-cysteine variants correlated with an increased likelihood of disulfide dimer formation, ii) N-terminal mutations were less likely to disrupt secretion, and were less prone to aggregation, iii) in contrast to wild-type FBLN3, multiple, but not all variants failed to induce MMP2 levels in cell culture, and iv) C-terminal mutations (either loss or gain of cysteines) were more prone to significant secretion defects, intracellular accumulation/misfolding, and stress response activation. These results provide molecular and biochemical insight into FBLN3 folding, secretion, and function for many cysteine mutations found in the human population, some of which may increase the likelihood of subclinical connective tissue or other FBLN3-associated haploinsufficiency diseases.

## INTRODUCTION

Extracellular matrix (ECM) proteins are critical regulators of cellular processes including cell communication, signaling, and adhesion (1). Accordingly, regulating the structure of ECM proteins is also similarly important for these processes. Mutations in ECM proteins (or proteins that in turn regulate ECM protein function) are associated with a series of diverse diseases ranging from connective tissue disorders like Marfan Syndrome (2, 3), marfanoid syndrome (4, 5), osteogenesis imperfecta (6, 7), to neurodegenerative diseases including age-related macular degeneration (AMD) (8–10) and retinal dystrophies (11–14).

Fibulin-3 (FBLN3, or EFEMP1) is a prime example an ECM protein wherein distinct mutations trigger diverse human diseases, including many of those mentioned previously. FBLN3 is a disulfide-rich ECM protein produced throughout the body and incorporated into basement membranes. The first identified mutation in FBLN3 was an autosomal dominant mutation (p.R345W) found in 1999 in a series of families with an AMD-resembling rare retinal dystrophy, Malattia Leventinese/Doyne Honeycomb Retinal Dystrophy (ML/DHRD) (14, 15). Since then, there has been a substantial growth in the number of FBLN3 mutations associated with disease. Collantes and colleagues identified multiple distinct autosomal dominant mutations (p.N80Y, p.R477C, p.494Glnext29) that cause juvenile-onset open angle glaucoma (16). Moreover, a homozygous recessive mutation (p.C55R) was the first reported loss-of-function (autosomal recessive) FBLN3 mutation associated with marfanoid symptoms (4, 5), but has since been joined by additional reports of mono-or biallelic FBLN3 mutations in individuals with connective tissue dysfunction (17–19). We speculated that additional rare FBLN3 mutations observed in the human population may also predispose FBLN3 to secretion defects, protein misfolding, and possibly stress response activation that ultimately may contribute to human disease.

In order to identify rare FBLN3 mutations, we turned to the ClinVar and gnomAD databases which provide genetic and mutational information extracted from deidentified individuals or population genetics. These resources can be used to begin to understand the genetic diversity of the human population at the gene level. Since FBLN3 is commonly included on exome sequencing panels for patients with presumed retinal dystrophies, it is not surprising that over 151 unique missense mutations in FBLN3 have been identified in these databases. Of these mutations, 24 involve cysteines (either gain-of-cysteine or loss-of-cysteine), which are residues known to be important structural and functional regulators of fibulin protein family members including FBLN3, fibulin-5 (FBLN5 or DANCE) (20–22), and EGF domains in general (which are canonical features of all fibulin proteins (23)). Thus, to better understand the complexity of disulfide bonding of FBLN3 and the consequences of rare cysteine mutations, we cloned ten new FBLN3 cysteine variants and evaluated their secretion, disulfide bonding, native molecular weight, effect on matrix metalloproteinase 2 (MMP2) activity, and endoplasmic reticulum stress (ER) stress response in culture. Our results demonstrate a surprising diversity of biochemical behaviors which may ultimately be leveraged to better understand FBLN3 and its relevance to a spectrum of human diseases.

## MATERIALS AND METHODS

### Plasmid generation

ClinVar (https://www.ncbi.nlm.nih.gov/clinvar/) and gnomAD (https://gnomad.broadinstitute.org/) were used to identify a series of seven cysteine mutations (Sup. Table 1) which originated from sequencing of clinical isolates. A previously identified C55R mutation (4, 5) along with five new loss-of-cysteine mutations (C42Y, C190R, C218R, C252F, C365S), two new gain-of-cysteine mutations (R358C, Y369C), and two additional random mutations (G57C, Y397C) were introduced into a pENTR1A 1xFLAG-tag (FT) WT FBLN3 DNA template (described previously (24)) using the Q5 Site-Directed Mutagenesis kit (New England Biolabs, Ipswich, ME). pENTRIA constructs were verified by Sanger sequencing before being introduced into the pcDNA DEST40 vector (Life Technologies, Carlsbad, CA) through an LR Clonase II reaction (Life Technologies, Carlsbad, CA). pcDNA constructs were also sequenced additionally prior to lipofection. Select mutations (G57C, C252F, and C365S) were introduced into a pENTR1A 3xFT HiBiT WT FBLN3 template, followed by recombination into the pLenti CMV Puro DEST vector (Addgene plasmid # 17452, gift from Eric Campeau and Paul Kaufman).

### Cell culture, transfections, and western blotting

For standard transfections, HEK-293A cells (R70507, Thermo Fisher Scientific, Waltham, MA) were grown in DMEM high glucose (4 g/L) media supplemented with 10% FBS and 1% PSQ (Gibco, Waltham, MA). The day before transfection, cells were plated at 75,000 cells per well of a 24 well plate (Corning, Corning, NY) and allowed to attach overnight. Cells were transfected with 500 ng pEGFP-N1 (for estimation of transfection efficiency) or the indicated DNA using Lipofectamine 3000/P3000 as described previously (24, 25). The following day, GFP-expressing cells were checked on a Celigo (Nexcelom, Lawrence, MA) for transfection efficiency (typically > 70%), and media was changed to serum free DMEM supplemented with 1% PSQ. Twenty-four hours after changing to serum free media, conditioned media was collected, and cells were washed and lysed using radio immunoprecipitation assay buffer (RIPA, Santa Cruz, Dallas, TX) supplemented with protease inhibitor (Pierce, Rockford, IL) and benzonase at 4°C for 10 min. Cell lysates were cleared by spinning 10 min at ≥ 20,000 x g in a chilled microcentrifuge. Soluble cell lysate was quantified using a bicinchoninic assay (Pierce) and normalized (typically to 20 μg). Samples were prepared with 1x SDS PAGE buffer with or without 2-mercaptoethanol (BME) and boiled for 5 min prior to loading on a 4-20% Tris-Gly gel. Gels were run for 90 min at 140V and transferred to nitrocellulose membranes using an iBlot2 (P0 program, Life Technologies). Membranes were stained with Ponceau S for total protein blocked overnight in PBS blocking buffer or Intercept blocking buffer (LI-COR). After washing in Tris-buffered saline supplemented with Tween-80 (0.5%, TBS-T), rabbit anti-flag (PA1-984B, 1:5,000, Pierce) and mouse anti-GAPDH (sc-47724, 1:10000, Santa Cruz) were added to membranes for 1 h in 5% BSA in TBS with NaN_3_ at RT, followed by goat anti-rabbit 680 (1:15,000, LI-COR) and goat anti-mouse 800 (1:15,000, LI-COR) in 5% milk in TBS-T for 40 min at RT. Blots were imaged on an Odyssey CLx and quantified using Image Studio software (LI-COR). A full blot image is shown in Supplemental Fig. 1A.

### Lentivirus production

To produce lentivirus, HEK-293T cells (CRL-3216, ATCC, Manassas, VA) were plated at 1.2 million cells/well of a 6 well plate (coated with poly-D-lysine to promote adherence). After allowing the cells to attach overnight, cells were transfected with psPAX2 (gift of Didier Trono, Addgene plasmid # 12260), VSvG (gift of Didier Trono, Addgene plasmid # 12259), Lipofectamine 3000, P3000, and each pLenti plasmid as described previously (5). The next day, media was changed and then collected at 24 and 48 h increments and pooled. Conditioned media containing lentivirus was spun down at 650 x g and filtered through a 0.45 μm PES filter before splitting into 500 μl single-use aliquots and freezing at −80°C.

### Stable cell line generation

ARPE-19 FBLN3 KO cells, described previously (5), were grown in DMEM/F12 supplemented with 10% FBS and 1% PSQ. These cells were used to avoid any influence on cell function by endogenous wild-type (WT) FBLN3. To generate stable cell lines expressing HiBiT-tagged FBLN3 variants, FBLN3 KO cells were plated at 300,000 cells/well in a 6 well plate. The next day, 500 μL of lentivirus supplemented with polybrene (1 μg/mL) was added to the cells for 24 h. Stable cells were selected by treating with 1 μg/mL puromycin at each media change for 1-2 weeks until no cell death was observed.

### HiBiT blotting

ARPE-19 FBLN3 KO cells expressing 3xFT HiBiT WT FBLN3, G57C, C252F, or C365S were plated on 24 well plates at 100,000 cells per well and incubated overnight. The next day, wells were changed to serum free media and incubated for an additional 48 h. Media and lysates were used for a HiBiT blot as described above for western blotting. After SDS-PAGE, separated proteins were transferred to nitrocellulose membranes (P0 program, iBlot2) which were washed with water followed by incubation in TBS-T for at least 30 min and incubation in HiBiT blotting buffer supplemented with LgBiT protein (Promega, Madison, WI) overnight at 4°C. The following day, blots were warmed to RT and NanoGlo luciferase substrate was added for 5 min prior to imaging on an Odyssey Fc (LI-COR). Subsequently, blots were blocked (Intercept, LI-COR) and probed for GAPDH as a loading control as described above. HiBiT blotting signal was quantified using Image Studio software (LI-COR). Full blot images are shown in Supplemental Fig. 1B, C.

### Size exclusion chromatography (SEC)

ARPE-19 cells (CRL-2302, ATCC) were electroporated with pcDNA or pLenti constructs encoding for 3xFT HiBiT WT FBLN3, G57C, C252F, or C365S. Briefly, 1 million cells were electroporated with 1 μg of midi-prepped plasmid DNA using a Neon NxT electroporation system (Life Technologies) with the following parameters: 1400V, 20 ms, x 2, similar to what we have described previously (26). Media was then changed to serum free media for 2-3 d. Conditioned media was collected and concentrated using a 30k molecular weight cutoff filter (Millipore, Burlington, MA). Samples were washed twice (buffer exchanged) with 10 mL of TBS. The final concentrate was then filtered through a 0.2 μm PES filter. Prior to loading onto a Superdex 200 Increase 10/300 column (Cytiva, Marlborough, MA) preequilibrated in TBS (0.25 mL/min) at 4°C. After loading, a 6 mL delay was used prior to collecting fractions every minute in a deep 96 well plate. Each fraction was then assayed for HiBiT activity by combining 25 μL of eluate with 25 μL of lytic HiBiT buffer supplemented with 1:100 LgBiT and 1:50 NanoBiT lytic substrate (27). Estimation of globular molecular weight was accomplished by running SEC Protein Standards (CellMosaic, Woburn, MA) followed by regression analysis.

### Transwell cell culture/zymography

ARPE19 FBLN3 KO and stable ARPE-19 HiBiT FBLN3 cells were plated at a density of 100,000 cells per well of a 12-well polyester transwell plate (0.4 μm pore, Corning) and media was changed to serum free the following day and 3-4 days thereafter. After two weeks (72 h since last media change), apical and basal media was collected for a HiBiT assay and zymography. Ten μL apical media and 20 μL basal media were combined with non-reducing SDS buffer and run on a 10% gelatin gel (Thermo Fisher Scientific) for 90 min at 140 V. Gels were incubated in renaturing buffer for 30 min at RT, then changed to developing buffer (G-Biosciences, St. Louis, MO) for an additional 30 min. Developing buffer was changed and gels were left gently shaking overnight at 37 °C. The following day the gels were stained with Coomassie R-250 for 1 h, then destained and rehydrated in water. Gels were imaged and analyzed on an Odyssey CLx using Image Studio software (LI-COR). Full zymography gels are presented in Supplemental Fig. 1D, E.

### Quantitative PCR (qPCR)

ARPE-19 FBLN3 KO and stable ARPE-19 HiBiT FBLN3 cells were plated on 12 well plates at 200,000 cells per well, allowed to attached overnight, followed by media change to serum free the next day. Forty-eight hours later, cells were trypsinized, pelleted, and washed with HBSS. Samples were stored at −80 until extraction using an Aurum Total RNA isolation kit (BioRad, Hercules, CA). mRNA was normalized and qScript cDNA SuperMix (Quantabio, Beverly, MA) was used to generate cDNA. Diluted cDNA was then amplified with TaqMan Advanced Fast master mix (Thermo Fisher Scientific) on a QuantStudio 6 (Thermo Fisher Scientific) using the following TaqMan probes (Thermo Fisher Scientific): hACTB (Hs01060665_g1), hDNAJB9 (Hs01052402_m1), hHSPA5 (Hs00607129_gH), hFBLN3 (Hs00244575_m1). Amplification results were normalized to ACTB, then to the 3xFT WT FBLN3 OE sample using the QuantStudio internal software.

## RESULTS

### Clinically-identified cysteine mutations in FBLN3 cause diverse biochemical behaviors

Disulfide bonding mediated through cysteine-cysteine covalent bonds are critical for the folding, secretion, and function of epidermal growth factor (EGF) domains (28, 29) and EGF-domain-containing proteins like FBLN3 (5, 21, 22). Accordingly, genetic mutation (either elimination or introduction) of critical cysteine residues is associated with FBLN3-related diseases, including marfanoid connective disease (4, 5) and juvenile open angle glaucoma (JOAG) (16). ClinVar (https://www.ncbi.nlm.nih.gov/clinvar/) and genomAD (https://gnomad.broadinstitute.org/) provide easily-accessible gene variant/mutation databases originating from genomic sequences from deidentified clinical isolates. To better understand how individual cysteines play critical roles in FBLN3, we tested five new loss-of-cysteine and four new gain-of-cysteine variants (Fig. 1A) in cell-culture-based biochemical and biophysical assays.

**Figure 1.**
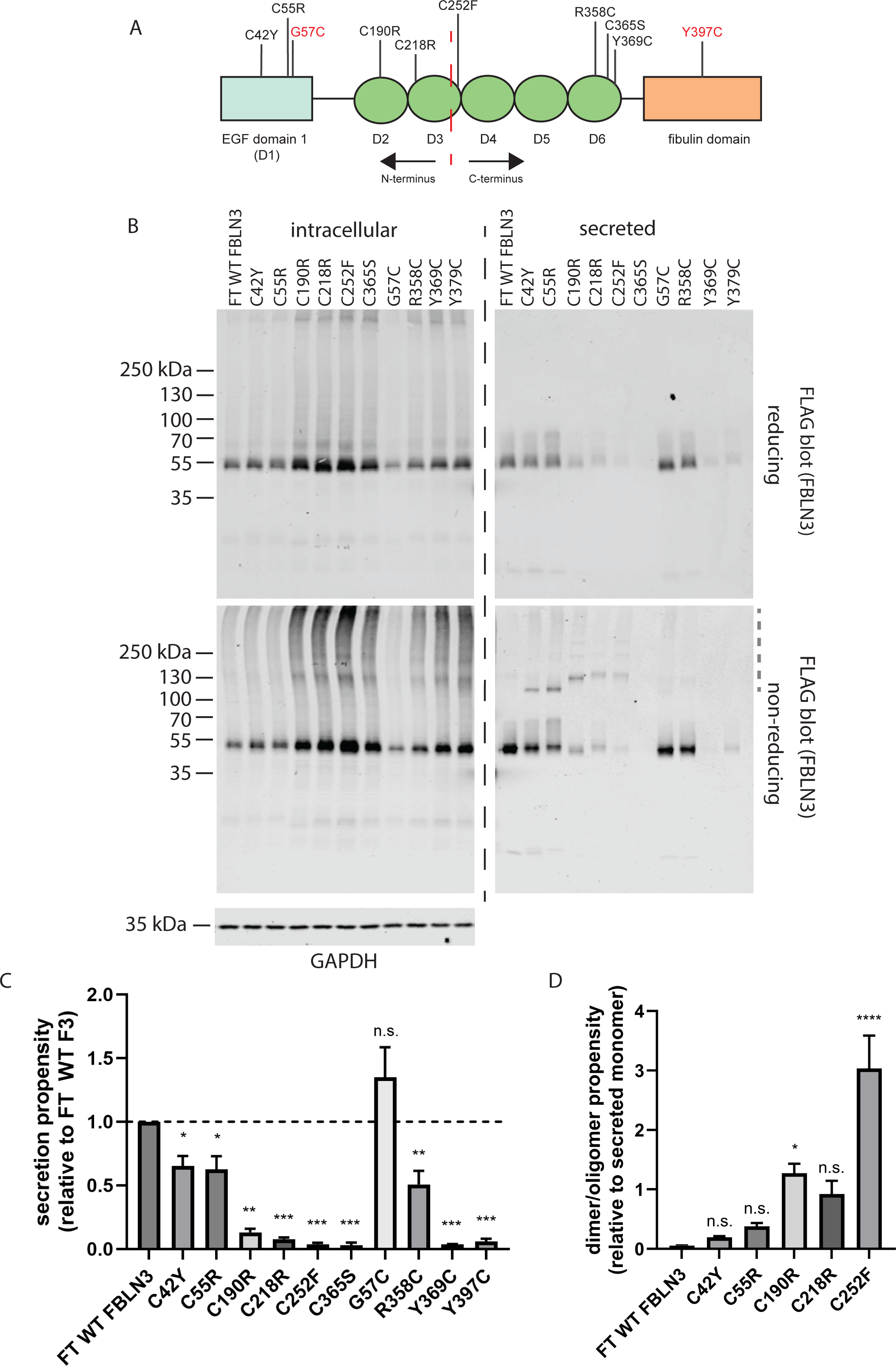
Clinically identified cysteine mutations affect FBLN3 secretion and disulfide bonding in diverse ways. (A) Cartoon schematic of FBLN3 domain structure and approximate location of each analyzed mutation. The red dashed line indicates an approximate half-way point (Asn249), providing N- and C-termini. Mutations shown in red indicate engineered variants. Light blue rectangle represents the atypical EGF domain whereas green ovals indicate conventional EGF domains. (B) HEK-293A cells were transfected with the indicated constructs followed by analysis of intracellular and secreted FBLN3 under reducing (top blots) and non-reducing (bottom blots) conditions. Representative images of at least three independent experiments. (C) Secretion propensity, defined as the quantification of secreted monomer/intracellular monomer under reducing conditions relative to WT FBLN3 was determined. n.s. = not significant, * p < 0.05, ** p < 0.01, *** p < 0.001, one sample t-test assuming a hypothetical mean of 1 (i.e., unchanged), mean ± S.D. (D) Dimer/oligomerization of FBLN3 is increased in select cysteine mutants. Disulfide dimers or oligomers (grey dashed line in non-reducing secreted FBLN3 blot) was normalized to monomer. * p < 0.05, **** p < 0.0001, one-way ANOVA vs. FT WT FBLN3 wtih Dunnett’s multiple comparisons test, mean ± S.D.

Consistent with our previous observations that C-terminal mutations tend to be more disruptive (less tolerated) than N-terminal ones (21, 24) with respect to FBLN3 secretion propensity, we again observed that mutations such as C42Y and C55R (studied previously) demonstrated higher secretion propensities (Fig. 1B, C, 0.65 ± 0.17 and 0.62 ± 0.23, respectively, relative to WT) than all other loss-of-cysteine mutations (C190R, C218R, C252F, C365S), which ranged from as low as 0.03 ± 0.03 (Fig. 1B-C, C365S) to 0.13 ± 0.05 (Fig. 1B-C, C190R) relative to WT FBLN3. Gain-of-cysteine mutations also followed a similar pattern wherein the N-terminal G57C mutation demonstrated no difference in secretion propensity relative to WT FBLN3 (Fig. 1B, C, 1.35 ± 0.62) contrasting all other analyzed gain-of-cysteine mutations (R358C, Y369C, Y397C), which ranged from 0.04 ± 0.003 (Fig. 1B, C, Y369C) to 0.51 ± 0.28 (Fig. 1B, C, R358C).

In addition to secretion propensity (defined as the secreted/intracellular ratio [reducing gel]), we also quantified the propensity for disulfide-linked dimer and oligomer formation in the conditioned media. Surprisingly, only the C190R and C252F mutations resulted in significantly higher disulfide-linked dimer/oligomer formation relative to WT FBLN3 (Fig. 1D), with C252F demonstrating the highest relative likelihood of higher molecular weight formation relative to monomer (Fig. 1D).

### FBLN3 cysteine mutant biochemical behavior reproduces in a retinal pigmented epithelial (RPE) cell line

While HEK-293A cells are a reasonable model to initially test a large number of FBLN3 variants, they are not a line that expresses high levels of endogenous FBLN3. Therefore, we also validated FBLN3 behavior in retinal pigmented epithelial (RPE) cells which produce high levels of FBLN3 (30, 31). Specifically, we tested how select cysteine variants behaved in ARPE-19 cells, an immortalized human RPE-derived line (32). Based on HEK-293A results, we selected i) G57C, a variant that was indistinguishable from WT (Fig. 1B, C), ii) C252F, a poorly secreted variant that demonstrated the highest propensity for dimer formation (Fig. 1D), and C365S, a variant with the highest propensity for intracellular retention (Fig. 1B, C). ARPE-19 FBLN3 knockout cells (FBLN3 KO, generated previously (5)) were used to avoid any confounding variables with endogenous WT FBLN3 during subsequent assays. Similar to our observations in HEK-293A cells, G57C behaved identically to WT FBLN3 with respect to secretion, intracellular levels and apparent disulfide bonding (Fig. 2A-C). Likewise, C252F and C365S both demonstrated secretion defects accompanied by intracellular accumulation that appeared to be a combination of disulfide-linked and covalent linkages (Fig. 2A-C). Interestingly, intracellular levels of cysteine mutations such as C252F and C365S appear to be more severe in ARPE-19 cells than in HEK-293A cells, suggesting that different cell types may handle FBLN3 mutants differently. Nonetheless, these results suggest that these particular mutations are not well tolerated across multiple cells of diverse origins.

**Figure 2.**
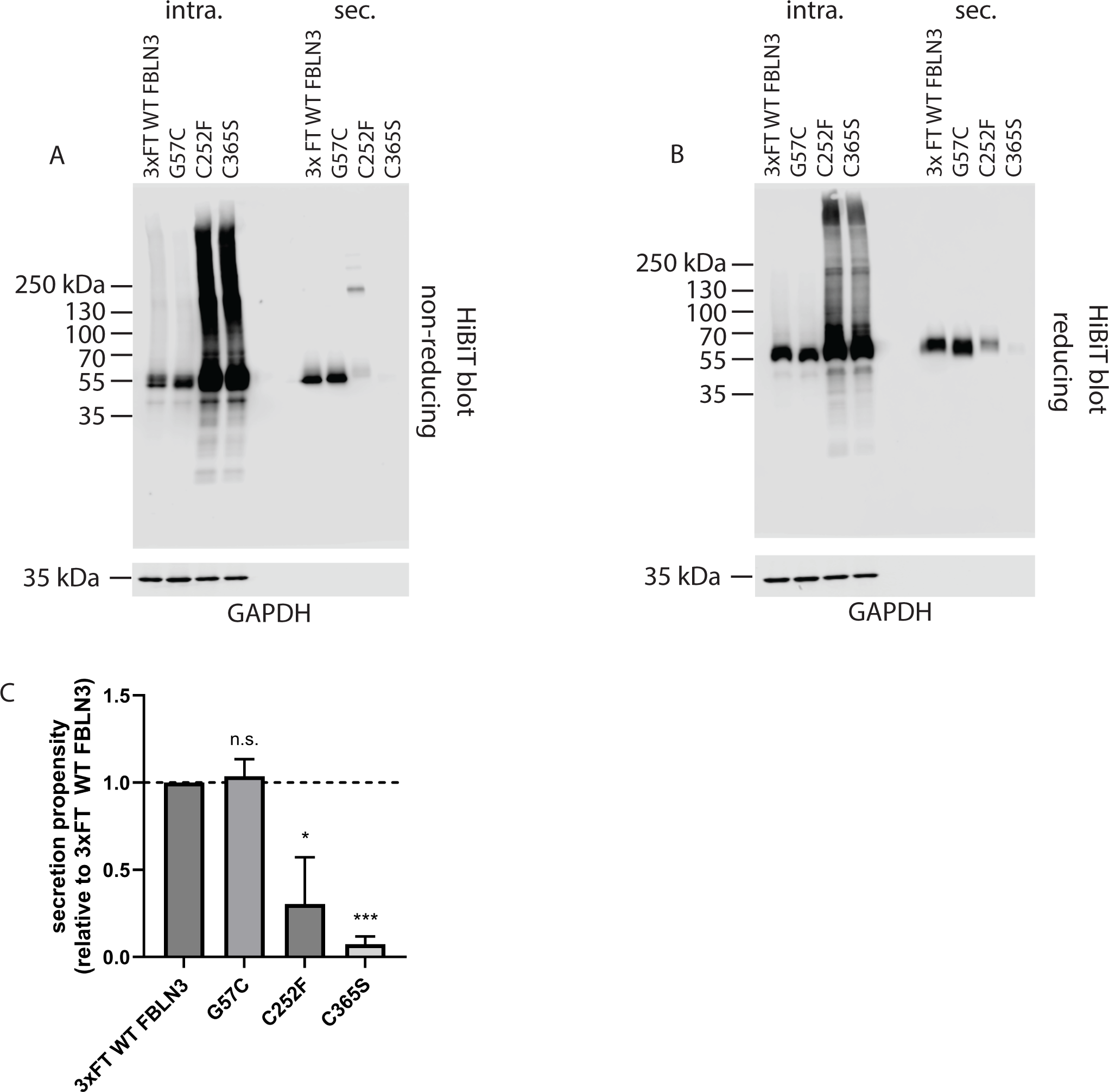
Cysteine mutant behavior reproduces in ARPE-19 cells. ARPE-19 cells were infected with lentivirus encoding for the indicated FBLN3 variant and stably selected. Intracellular and secreted FBLN3 were evaluated under non-reducing (A) and reducing (B) conditions. Representative images of three independent experiments. (C) Secretion propensity of the apparent FBLN3 monomer under reducing conditions was calculated, mean ± S.E.M., n.s. = not significant, * p < 0.05, *** p 0.001, one sample t-test vs. hypothetical mean of 1 (i.e., unchanged).

### Size exclusion chromatography (SEC) suggests that FBLN3 migrates as an apparent dimer under native conditions

Most analyses of FBLN3 have been performed under denaturing SDS-PAGE conditions. To determine whether select cysteine mutations affected FBLN3 native confirmation or quaternary structure, we developed a technique that pairs apparent native size resolution (SEC) with the sensitivity of a HiBiT assay. We were surprised to find that secreted WT FBLN3 migrates primarily as a ∼110 kDa species (Fig. 3A), that is equivalent to a FBLN3 dimer based on an estimated monomeric molecular weight of 54.6 kDa (https://www.uniprot.org/uniprotkb/Q12805). An additional WT FBLN3 species eluted after the void fraction till ∼11 mL, encompassing a poorly defined range of species between ∼669 kDa and ∼232 kDa (Fig. 3A). Perhaps even more interestingly was that secreted versions of G57C and C365S both migrated identically to WT FBLN3 (Fig. 3B, D), although the absolute HiBiT signal in both C252F and C365S samples were substantially reduced, as demonstrated in Fig. 1 and Fig. 2. Migration of the main C252F species was slightly altered compared to the other three FBLN3 variants, peaking at ∼122 kDa (Fig. 3C). Additional higher molecular weight C252F species were observed at ∼231 kDa, equivalent to a species resembling a dimer of dimers, and ∼602 kDa, which may correspond to soluble, but aggregated protein, or perhaps an octamer (two dimer-dimers) (Fig. 3C). Overall, these observations provide corroboration of our SDS PAGE results, but also add an additional dimension to FBLN3 that will need to be further explored and defined for additional FBLN3 variants.

**Figure 3.**
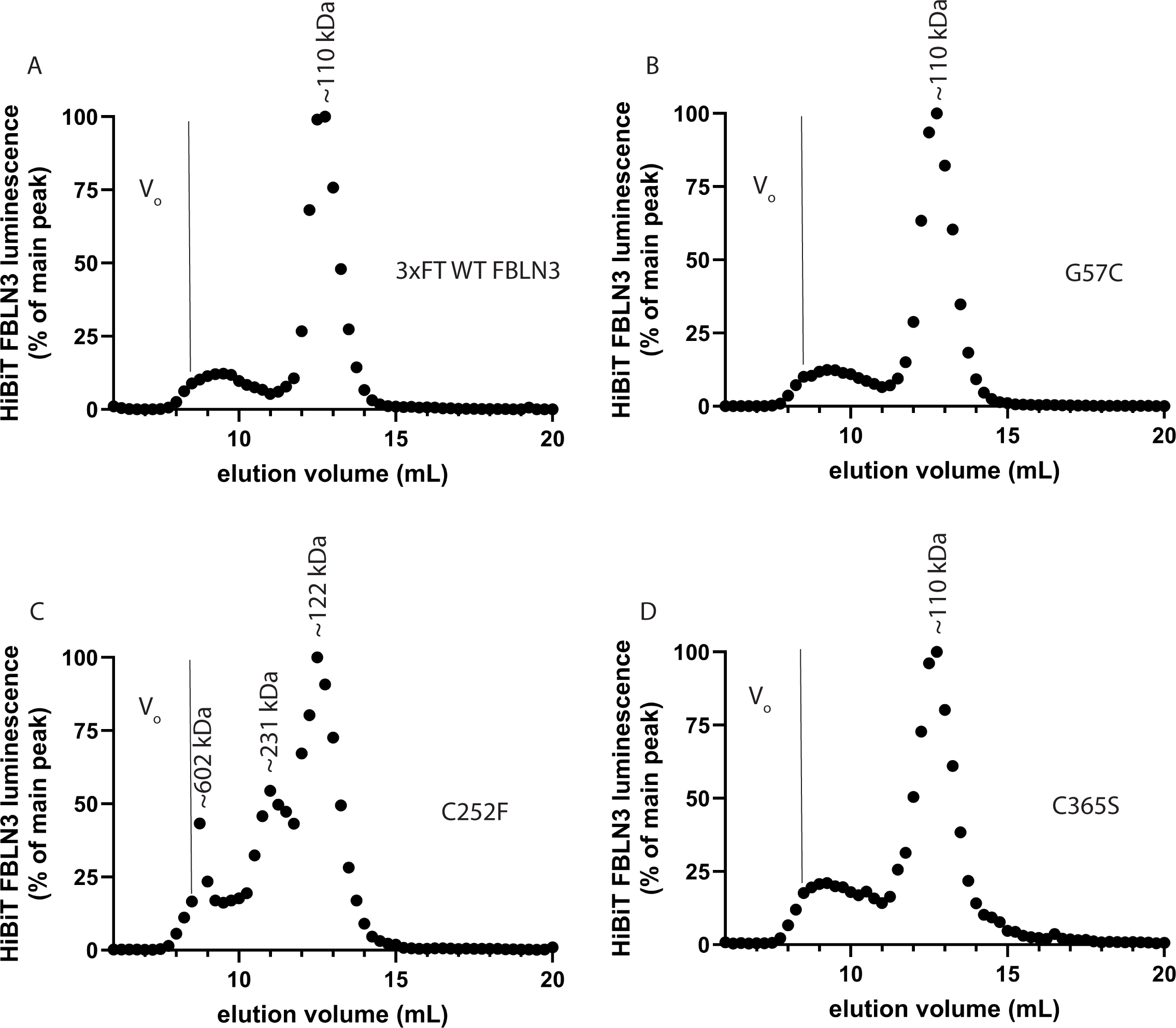
Size exclusion chromatography (SEC) demonstrates native secreted FBLN3 apparent molecular weight and quaternary structure. ARPE-19 cells were electroporated with constructs encoding for 3xFT HiBiT FBLN3 followed by exchanging media into serum free conditions. After concentration of the conditioned media using a 30,000 molecular weight cut off filter and washing twice in TBS, samples were loaded onto a Superdex 200 Increase 10/300 also equilibrated with TBS. Fractions (0.25 mL) were collected every min after a 6 mL delay and subjected to a HiBiT assay to generate a ‘chromatogram’. Representative chromatograms of two independent experiments. Indicated apparent molecular weights are based on interpolation of a standard curve using standards (CellMosaic). V_o_ = void volume, >669 kDa.

### Inefficiently secreted FBLN3 cysteine variants are incapable of regulating MMP2 levels

While FBLN3 has no well-accepted primary function, previous cell culture studies have demonstrated that WT FBLN3 increases MMP2 activity (5, 33) as determined by zymography. To obtain a better sense of whether select cysteine variants could behave functionally similar to WT FBLN3 in this respect, we assessed apical and basal MMP2 levels in stable ARPE-19 cells expressing WT, G57C, C252F, or C365S FBLN3 that also lack endogenous FBLN3 (FBLN3 KO). Only WT and G57C FBLN3 expressing cells demonstrated a significantly higher level of MMP2 in both the apical and basal chambers compared to FBLN3 KO cells (Fig. 4A, B) indicating functional FBLN3. In fact, we found that G57C, which has indistinguishable secretion and native migration parameters compared to WT FBLN3, significantly increased apical and basal MMP2 levels even relative to WT FBLN3 (Fig. 4A, B). Conversely, both C252F and C365S failed to elevate apical and basal MMP2 levels (Fig. 4A, B), suggesting that they may lack the inherent ability to elevate MMP2, or that their secreted levels are too low to alter MMP2 at a detectable level.

**Figure 4.**
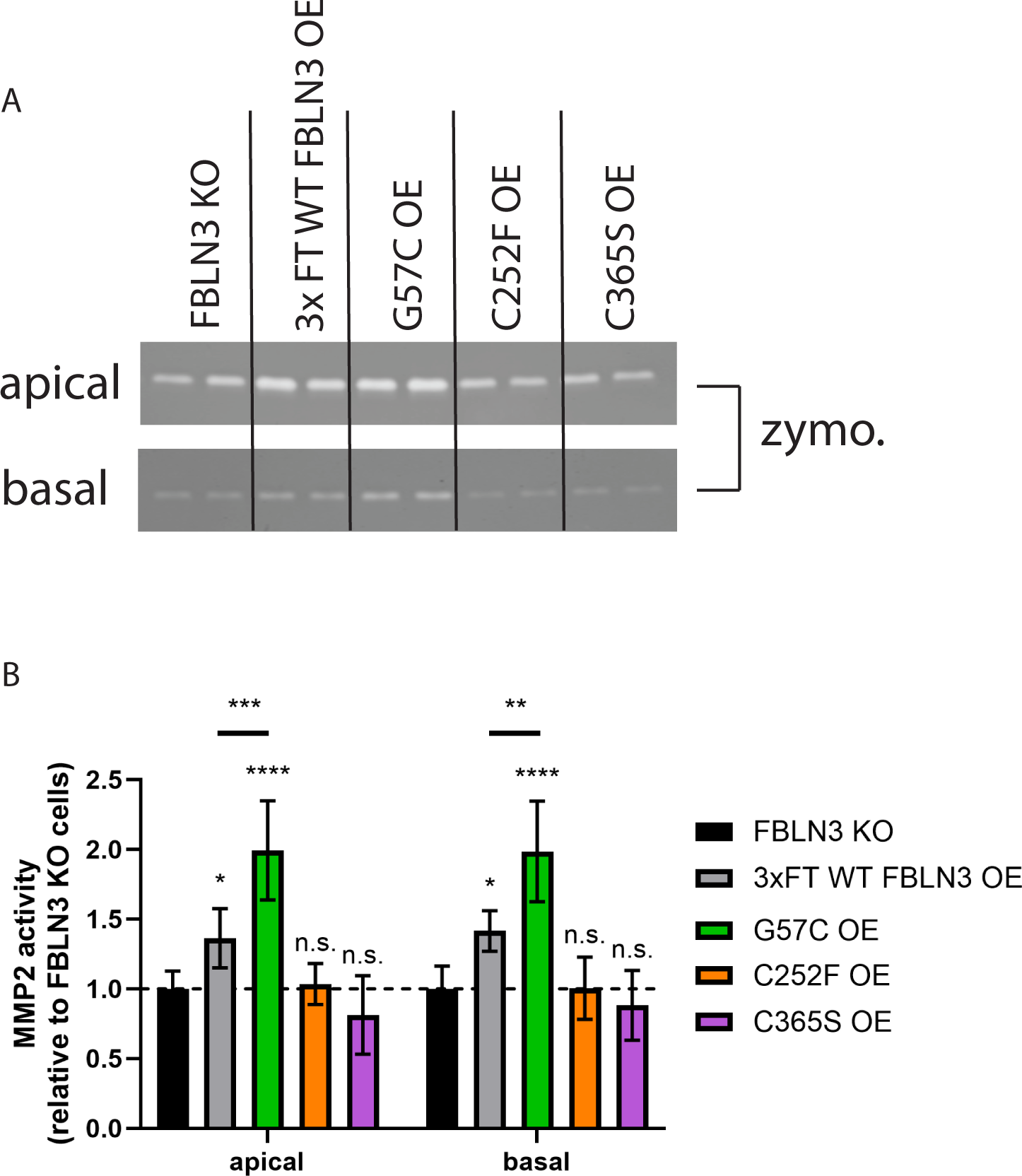
FBLN3 cysteine mutations affect MMP2 zymography-based activities. (A) Stable ARPE-19 cells (lacking endogenous FBLN3, “FBLN3 KO”, same as those described in Fig. 2) were plated on 0.4 μm 12 well transwells followed by switching media to serum free conditions for 2 wks. Apical and basal aliquots of media were taken after 72 h of accumulation and analyzed on a 10% gelatin gel. (B) Quantification of MMP2 activity in different stable cell lines. n = 3 independent experiments, mean ± S.D. n.s. = not significant, * p < 0.05, ** p 0.01, *** p < 0.001, one-way ANOVA vs. FBLN3 KO cells with Dunnett’s multiple comparisons test. An additional one-way ANOVA was performed comparing 3xFT WT FBLN3 OE to G57C OE.

### Intracellular FBLN3 triggers endoplasmic reticulum (ER) stress response activation

Given the high levels of intracellular FBLN3 observed in inefficiently secreted variants (Fig. 1B, Fig. 2A, B), and the characteristic smeared intracellular banding pattern commonly associated with aggregated, misfolded proteins, we tested whether ER stress was triggered in select cysteine variants chosen for follow up studies. As expected, levels of FBLN3 in KO cells were significantly lower than in all cell lines with overexpressed FBLN3, and, importantly, the levels of overexpressed FBLN3 were not different across WT, G57C, C252F, or C365S lines (Fig. 5). Consistent with our previous observations (22), overexpression of WT FBLN3 did not trigger ER stress response activation as indicated by transcriptional levels of DNAJB9 (indicative of IRE1 signaling (34, 35)) or HSPA5 (indicative of ATF6 signaling (34, 35)) (Fig. 5). However, we did detect a significant increase in both DNAJB9 and HSPA5 levels triggered by expression of both C252F and C365S (Fig. 5), indicating that indeed these variants are likely misfolded intracellularly in addition to being incapable of exerting FBLN3-dependent effects extracellularly as demonstrated in the MMP2 zymography experiments. These results suggest that FBLN3 cysteine mutations may have detrimental physiologic effects (i.e., stress response activation) while also potentially exerting a FBLN3 haploinsufficiency in the individuals who harbor these mutations.

**Figure 5.**
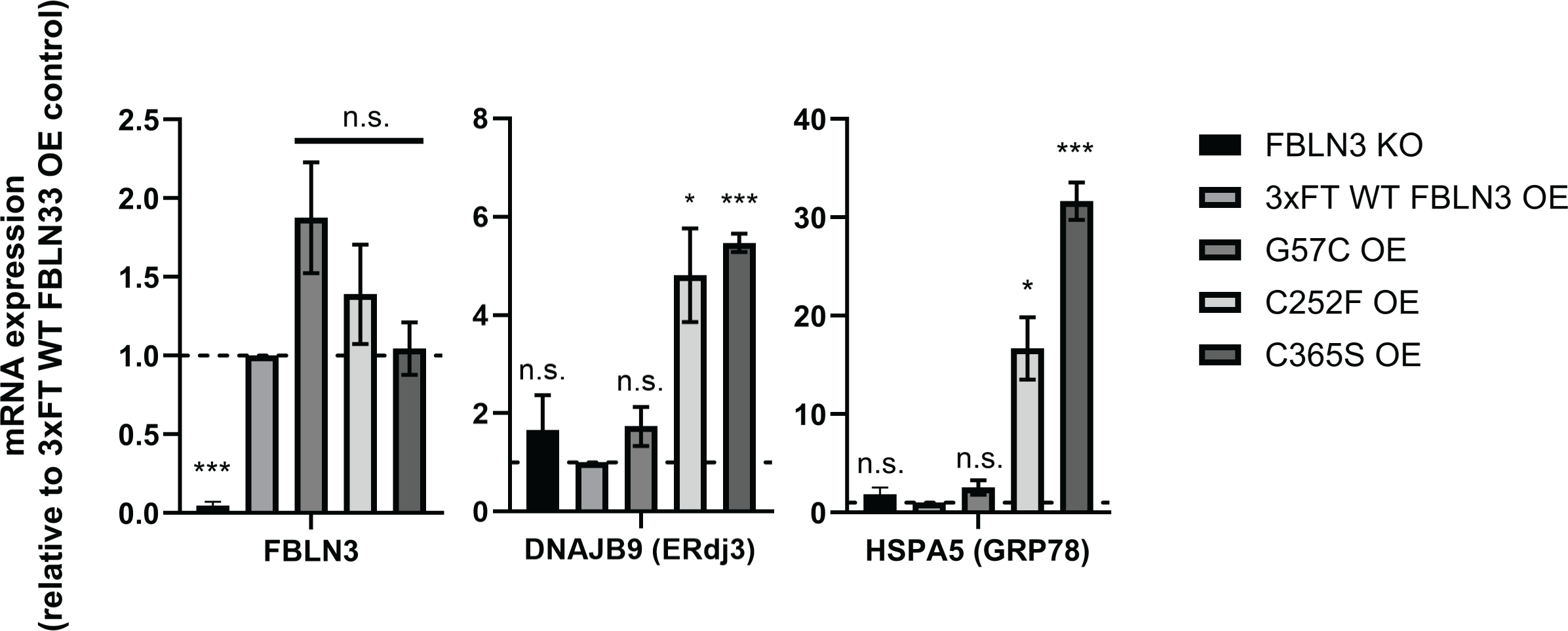
Inefficiently secreted FBLN3 cysteine variants trigger increased endoplasmic reticulum (ER) stress. Stable ARPE-19 cells were analyzed by quantitative PCR (qPCR) for FBLN3, DNAJB9 (ERdj3), and HSPA5 (GRP78) transcript levels. Three to four independent data sets were combined for each panel, mean ± S.E.M, n.s. = not significant, * p < 0.05, ** p 0.01, *** p < 0.001, one sample t-test vs. hypothetical mean of 1 (i.e., unchanged).

## DISCUSSION

FBLN3 is increasingly becoming accepted as an intriguing and dynamic ECM protein important to human health and disease (36). As evidence of this, mutations in FBLN3 have been associated with diverse diseases ranging from connective tissue disorders (4, 17–19) to blinding eye diseases (14, 16, 31, 37). Previously we characterized a series of clinically-identified missense mutations across FBLN3, identifying particular variants that were prone to aggregation and potentially associated with rare forms of retinal degeneration (24). Herein, we have expanded upon these studies, including seven cysteine variants pulled from online clinical and population databases along with two engineered cysteine variants. While all the mutations that we tested would be considered rare to ultra-rare with an allele count (# of chromosomes sequenced) ranging from 1 (C218R, C252F, C365S) to 12 (R358C) (Sup. Table 1), the data extracted from these unique variants can be used to provide a better understanding of FBLN3 folding, secretion, and potential involvement in pathology.

We speculate that distinct biochemical behaviors may dictate the ultimate FBLN3-based disease to which variants potentially contribute. For example, genetic (or functional) loss of one or more copies of FBLN3 correlates with association to connective tissue diseases such as Marfan-like or marfanoid syndrome. While the majority of mutations associated with this spectrum of FBLN3-diseases are caused by non-sense mutations or premature stop codons leading to non-sense mediated mRNA decay (17–19), there are exceptions to this rule, most notably a homozygous p.C55R mutation (4). This mutation results in FBLN3 protein production and secretion, but also causes the formation of aberrant extracellular disulfide bonds and results in a non-functional FBLN3 protein incapable of regulating MMP2 levels or aligning collagen IV fibers (5). Conversely, mutations that appear to cause aberrant intracellular and/or extracellular FBLN3 protein misfolding correlate with either retinal degeneration (13, 14, 37) or glaucoma (16, 38). While we (22) and others (13) have demonstrated that certain FBLN3 mutations likely don’t elicit an ER stress response, we cannot rule out that additional, newly-identified FBLN3 mutations such as those involved in glaucoma, do in fact trigger ER stress. These gaps in knowledge highlight how little we still know about how FBLN3 mutations can cause distinct ocular disease like retinal degeneration but not glaucoma, and vice versa. Many of the mutations that we have characterized in this work would seem to fall somewhere in the grey zone of being likely haploinsufficient due to secretion defects, but also aggregation-prone as demonstrated by western/HiBiT blotting and through the activation of stress-responsive signaling pathways in the ER via qPCR. Aside from the R358C mutation originating from an “inborn genetic disease” condition, the ClinVar database did not provide any associated phenotypes any of the cysteine mutations, and all are considered variants of unknown significance that require additional genetic and/or functional input for more definitive classification.

Our findings also highlight a series of underappreciated properties of FBLN3. For example, in our previous studies, we engineered a series of cysteine mutations that had distinct extracellular disulfide bonding properties depending on where the cysteine mutation occurred (5). At the time, we were not sure if these same bonding patterns happened in humans, but upon following up in these studies, we find that they in fact do occur. Based on reducing and non-reducing SDS PAGE analysis, cysteine mutants of FBLN3 clearly have the ability to dimerize, as we first demonstrated with the C55R mutation, but our current SEC studies suggest that even WT FBLN3 appears to also be capable of migrating as an apparent non-covalent dimer under native conditions. In fact, this is the main species present for WT FBLN3 under native conditions. Moreover, SEC analysis also emphasizes the ability of FBLN3 to form aggregates or oligomers, which are further aggravated by select FBLN3 mutations, like C252F, and possibly additional mutations. Yet, what these species mean from a biological or pathophysiological perspective is unclear. Could they be toxic soluble oligomers (39, 40), or the seeds of amyloid formation (41, 42)? More definitive biochemical studies are required to test these possibilities. Nonetheless, overall, our findings establish the basis for beginning to unravel the complexity of FBLN3 folding, disulfide bond formation, and the triggering of ER stress, all of which may play a part in the role of FBLN3 in regulating human health and disease.

## ACKNOWLEDGEMENTS

JDH is the Larson Endowed Chair for Macular Degeneration Research (UMN). JDH is supported by R01-EY027785.

