## Supplemental Fig for "A spectrum of clinically-identified cysteine mutations in fibulin-3 (EFEMP1) highlight its disulfide bonding complexity and potential to induce stress response activation"

Sup. Fig. 1

A

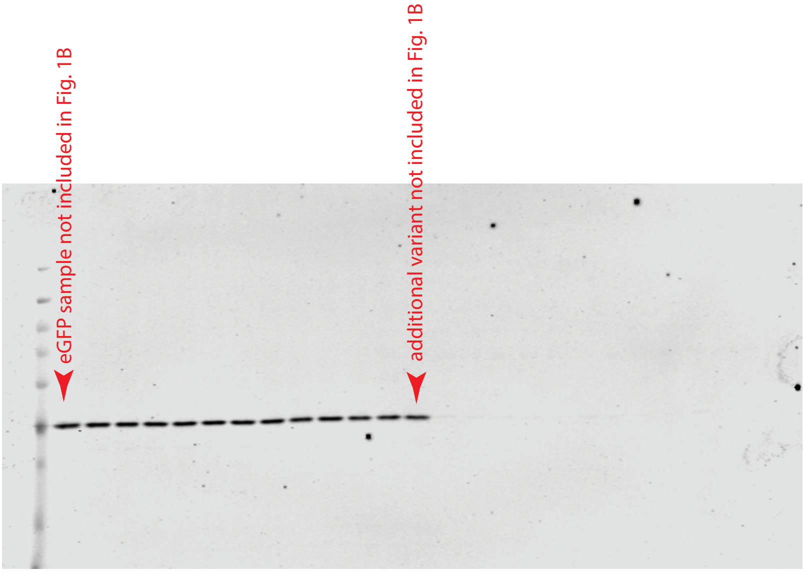


full blot used for GAPDH in Fig. 1B

B

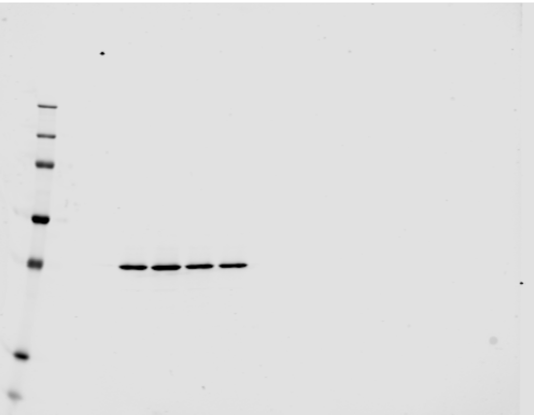

full blot used for GAPDH in Fig. 2A

C

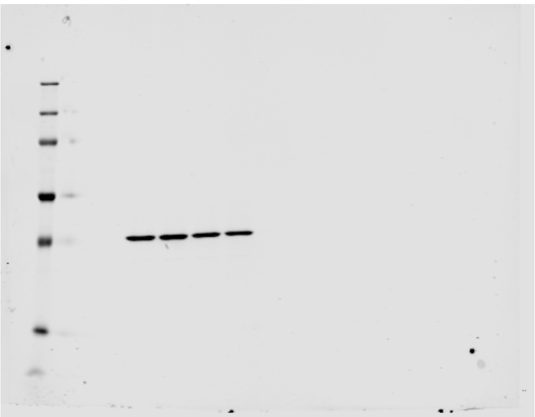

full blot used for GAPDH in Fig. 2B

D

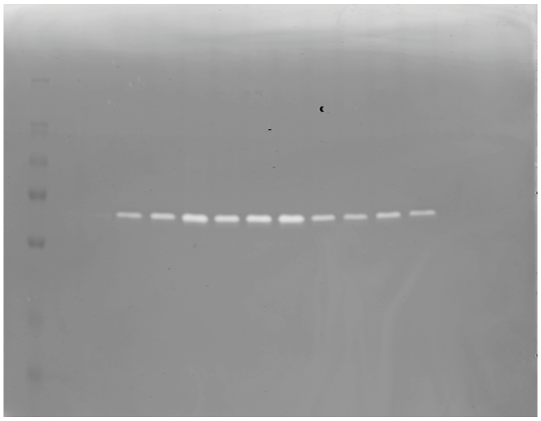

full gel used for zymography in Fig. 2D

E

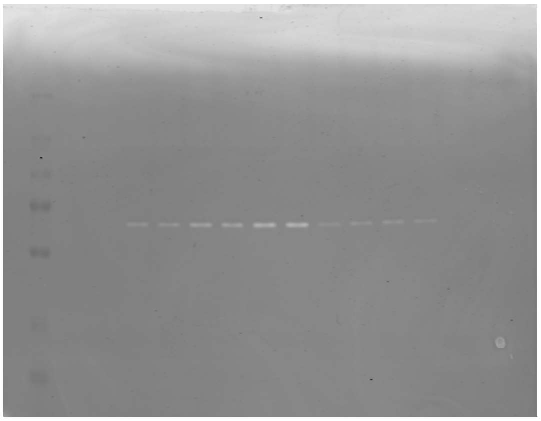

full gel used for zymography in Fig. 2E
