## Supplemental Table for "A spectrum of clinically-identified cysteine mutations in fibulin-3 (EFEMP1) highlight its disulfide bonding complexity and potential to induce stress response activation"

Sup. Table 1

| gnomAD ID | Protein Consequence | Transcript Consequence | ClinVar Clinical Significance | ClinVar Variation ID | Allele Count | Allele Frequency |
|---|---|---|---|---|---|---|
| 2-55918224-C-T | p.Cys42Tyr | c.125G>A | | | 4 | 2.48E-06 |
| 2-55881684-A-G | p.Cys190Arg | c.568T>C | Uncertain significance | 2006112 | 6 | 3.72E-06 |
| 2-55877854-A-G | p.Cys218Arg | c.652T>C | | | 1 | 6.20E-07 |
| 2-55877751-C-A | p.Cys252Phe | c.755G>T | | | 1 | 6.20E-07 |
| 2-55871030-C-G | p.Cys365Ser | c.1094G>C | | | 1 | 6.20E-07 |
| 2-55871052-G-A | p.Arg358Cys | c.1072C>T | Uncertain significance | 2259874 | 12 | 7.44E-06 |
| 2-55871018-T-C | p.Tyr369Cys | c.1106A>G | | | 2 | 1.24E-06 |
